# Decoding the functional Fes kinase signaling network topology in a lymphocyte model

**DOI:** 10.1101/125088

**Authors:** Andreas O. Helbig, Michael Kofler, Gerald Gish, Lourdes Sriraja, Kristina Lorenzen, Monika Tucholska, Cunjie Zhang, Frederick P. Roth, Tony Pawson, Karen Colwill, Evangelia Petsalaki

## Abstract

The c-Fes protein tyrosine kinase is a proto-oncogene that can also act as a tumor suppressor. We implemented an unbiased phosphoproteomics-based analysis that establishes cognate kinase-substrate associations, and revealed that c-Fes directly phosphorylates Dok1, Ptpn18 and Sts1, facilitating recruitment of the Src inhibitory kinase Csk to these substrates. These interactions resulted in modulation of Src signaling following B-cell receptor (BCR) stimulation and subsequent alteration of the protein levels of CD19, a membrane-localized BCR co-receptor and emerging key protein affecting the development of B- and plasma cell-lymphoma. Strikingly, manipulating c-Fes expression levels drove opposing biological outcomes. Low-level exogenous c-Fes expression led to a strong increase in CD19 protein levels while high c-Fes expression abolished CD19 protein levels. Thus, we propose that a balance of c-Fes and Src signaling can regulate receptor maintenance, which may influence cellular outcome such as tumorigenesis or tumor suppression.

## Introduction

Over the last 30 years, studies on the cytoplasmic tyrosine kinase Fes and the related kinase Fer have provided key insights into kinase structure and general protein domain architecture, particularly the discovery of the non-catalytic phosphotyrosine binding SH2 domain (Sadowski et al., 1986), (DeClue et al., 1987). Fes and Fer share a unique domain composition: An N-terminal F-BAR domain, a central SH2 domain, and a C-terminal tyrosine kinase domain. F-BAR domains interact with phosphoinositides to promote membrane localization and invagination (Tsujita et al., 2006), (McPherson et al., 2009), while the phosphotyrosine binding SH2-domain localizes and activates the catalytic domain of soluble tyrosine kinases, thereby processing and propagating intracellular signals. Although biological functions have been ascribed to its individual domains, the molecular basis for Fes’s biological functions is still poorly understood. Only a few Fes substrates such as STATs, CTTN and Bcr have been identified so far (Greer, 2002), and little is known about how this kinase’s phosphorylation events influence cellular function.

c-Fes knockout macrophages display immune hypersensitivity which points to Fes being involved in the regulation of the immune response (Parsons and Greer, 2006). Indeed, Fes is highly expressed in the myeloid system where it induces the differentiation of myeloid progenitor cells into terminal macrophages (Carè et al., 1994), (Kim and Feldman, 2002). It is also expressed in several other lymphoma lines and blood tissues (MacDonald et al., 1985) suggesting that Fes is active in multiple lineages of the hematopoietic system. Fes, along with Fer, is activated downstream of oncogenic KIT receptors in primary acute myeloid leukemia (AML) and is expressed in AML cell lines, where it is necessary for cell proliferation and survival (Voisset et al., 2010). This activating role of Fes in AML contrasts with its apparent role as a tumor suppressor in colorectal cancer (Bardelli, 2003), (Sangrar et al., 2005), and underscores the need for a better understanding of Fes-modulated signaling networks. Particularly, a detailed grasp of *in-vivo* substrates is necessary to link such observations to a molecular basis. Therefore, a data-driven systems biology approach, providing comprehensive information on direct substrates and their operation within the signaling network is crucial to place Fes into a more defined functional context. Here, we employed a quantitative phosphoproteomics and bioinformatics approach to elucidate the mechanistic context of Fes kinase function and in parallel provide a systemic resource of kinase-substrate associations in B-lymphocytes.

## Results

### Fes activation and selectivity in B-cells

To better understand the signaling pathways influenced by the Fes kinase, we sought a biological process where catalytic activity could be induced by the application of a biological stimulus. Focusing on the hematopoietic system, where there is evidence for Fes activity (Scheijen and Griffin, 2002), we tested the expression of Fes in the RAW 264.7 macrophage cell line as a representative of the myeloid system and in the human B-lymphocyte cell line DG75 as a model for the lymphoid system. Since the abundance of Fes in DG75 cells was relatively low compared to RAW 264.7 macrophages (Figure S1A), we chose to conduct further experiments in DG75 cells. Here, we could effectively alter Fes signaling by exogenous Fes kinase expression, a strategy that would be much less effective in myeloid model systems already containing a high baseline of endogenous Fes signaling. Immunofluorescence microscopy indicated that endogenous Fes co-localized with the B-cell Receptor (BCR) in resting DG75 cells (Figure 1A). Further analysis to evaluate the model system indicated that DG75 cells indeed bear hallmarks of B-cell signaling including expression of B-cell markers and activation of downstream signaling pathway components after IgM stimulation (Figure 1B, Figure S1B-C). BCR stimulation through antibody cross-linking reproducibly led to endogenous Fes phosphorylation on its activating site, Y713 (Figure 1C). As the small molecule Fes inhibitor TAE684 (Hellwig et al., 2012) effectively inhibited the pY713 signal, the activation process appeared to involve auto-phosphorylation and not the action of another kinase working in trans. Similarly, BCR stimulation activated stably expressed GFP-tagged full-length Fes or a catalytically active SH2-kinase fragment in DG75 cells as measured by Y713 auto-phosphorylation (Figure 1D). Curiously, a stronger pY713 signal was observed for the full-length F-BAR domain-containing protein than the truncated Fes derivative following stimulation in cells. To determine the extent by which the F-BAR domain contributes to kinase function, we performed an *in-vitro* substrate phosphorylation analysis by applying a broad-spectrum kinase inhibitor (FSBA) to DG75 cell lysate then incubating the kinase-dead lysate with purified full-length Fes or SH2-kinase. The added kinase amounts were normalized on kinase activity of the recombinant proteins. As detected by an anti-phosphotyrosine antibody (4G10) immunoblot, the exogenous Fes kinases were observed at the expected molecular weight (Figure 1E). The number of phosphorylated target proteins was lower for the F-BAR containing full-length Fes even though the level of activity as indicated by auto-phosphorylation level for full length and truncated kinases was comparable. This indicates that the F-BAR domain plays a unique part in establishing substrate specificity (Figure 1E) as well as enabling kinase activation following BCR stimulation (Figure 1D).

**Figure 1.**
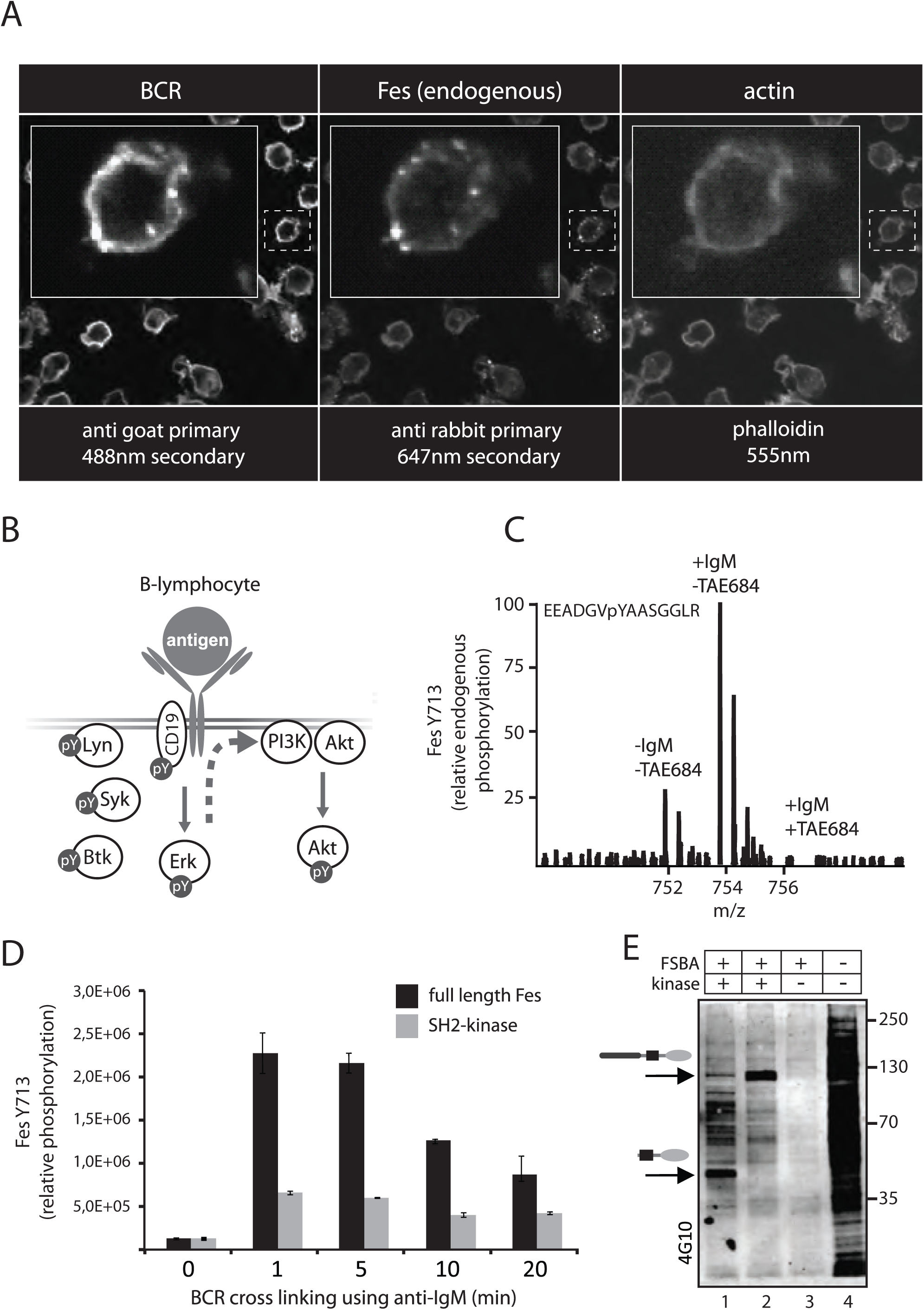
The Fes kinase is an active signaling component in the DG75 B-lymphocyte cell line. (A) Immunofluorescence microscopy showing co-localization of endogenous Fes kinase with the B-cell receptor in fixed DG75 cells. (B) Schematic representation of the engagement of major B-cell receptor (BCR) signaling components, following receptor clustering after antigen binding. Src family kinase Lyn becomes activated recruiting downstream tyrosine kinases such as Syk to receptor signaling complexes. Following this, there is an activation of the MAP kinase cascade as well as recruitment of PI3K to the membrane and activation of Akt signaling. (C) Phosphoproteomic analysis of DG75 lysates showed that endogenous Fes kinase autophosphorylation on Y713 is increased following BCR stimulation for 10 minutes (+IgM). Intensities of the auto phosphorylated peptide (pY713) were normalized to five unmodified peptides from the SH2 and kinase domain. Samples were compared using dimethyl labeling. The compound TAE684 (1uM final concentration) was used to inhibit Fes kinase activity. (D) Multiple reaction monitoring (MRM) mass spectrometry analysis of the Fes auto-phosphorylation site during BCR cross linking. DG75 cells were stably transfected with GFP-tagged full length Fes or an SH2-kinase version lacking the F-BAR domain. Both constructs were expressed at similar levels. Intensities of the auto-phosphorylated peptide (pY713) were normalized to five unmodified peptides from the SH2 and kinase domain. (E) Western blot analysis, using an anti-phosphotyrosine antibody (4G10), of kinase assays on whole cell lysates was performed. DG75 cell lysate was rendered kinase-dead in advance by a broad spectrum kinase inhibitor (FSBA) treatment. Following this, equal amounts of Fes kinase activity were added for the respective kinases as measured by amount of kinase autophosphorylation (indicated by arrow). In lane 1, purified Fes SH2-kinase was added. Lane 2 was treated with the full-length Fes protein. Lane 3 contained inhibited lysate without added kinase. Lane 4 contained uninhibited cell lysate subjected to kinase assay conditions.

### Fes signaling within the B-cell phosphotyrosine network

To analyze kinase signaling downstream of BCR activation, a phosphoproteomics-centered strategy was implemented. We manipulated Fes activity through stable or transient expression of c-Fes or an activated Fes kinase E708A (Figure S2) to define direct Fes substrates and highlight its influence within the global signaling network (Figure 2A). The BCR of DG75 cells was then activated using anti-human IgM to establish a global picture of BCR-related signaling events also with Fes modulation. Inhibition of endogenous Fes signaling using TAE684 was complemented with other kinase inhibitors (PP2, SrcI1, PD184352) to establish signaling direction and topology. We employed a protocol that maximized phosphoproteome capture through TiO_2_ enrichment of phosphorylated tryptic peptides followed by strong cation exchange (SCX) chromatography or immunoprecipitation of tyrosine phosphorylated peptides prior to mass spectrometry analysis. Dimethyl chemical labeling was utilized to compare three individual samples in one MS experiment (Figure 2B).

**Figure 2.**
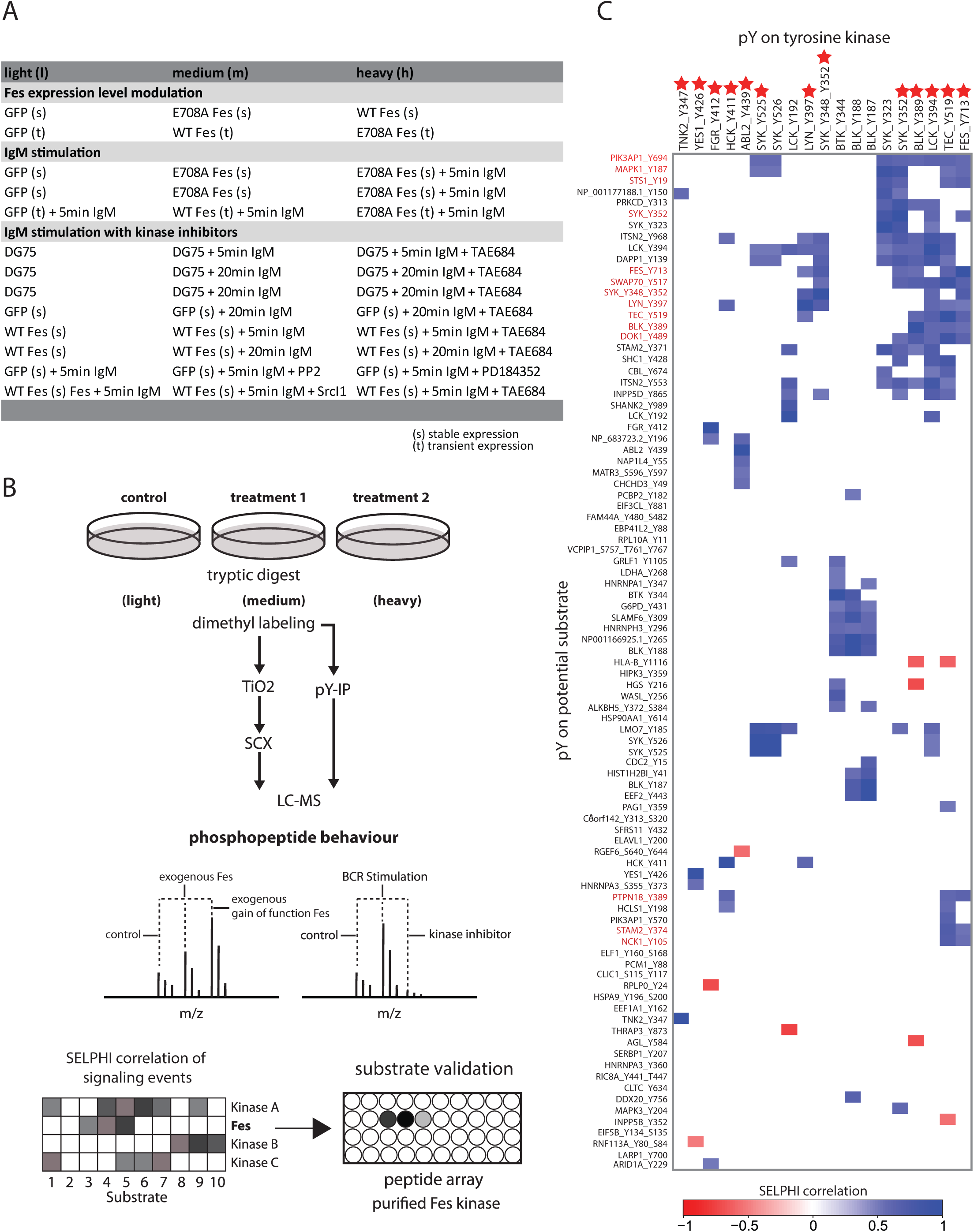
Analytical strategy reveals global signaling events and kinase substrate associations. (A) Table of phosphoproteomic experiments performed on the DG75 lymphocyte cell line. Kinase signaling pathways were perturbed by transient expression (t) or stable integration (s) of exogenous wildtype Fes kinase as well as a gain of function kinase variant (E708A). Cells were stimulated by IgM cross linking for 5 or 20 minutes as indicated. Additional large-scale experiments evaluated the impact of Fes, Src and Mek inhibitors on signaling events. (B) After tryptic digestion and C18 peptide purification, peptides were labeled using chemical dimethyl labeling and phosphorylated peptides were isolated using TiO2 enrichment as well as anti-phosphotyrosine immunoprecipitation. TiO2 eluates were subsequently fractionated using strong cation exchange chromatography. Mass spectrometric analysis using an Orbitrap elite was followed by Maxquant data analysis (Table S1). Figure shows example of conditions tested. Data integration and correlative analysis was performed using the SELPHI platform. This procedure allowed us to construct maps for potential kinase-substrate associations (Table S2/S3). In particular, tyrosine phosphorylation events were used to build peptide arrays that were subsequently probed with purified Fes kinase to validate the bioinformatics results (Table S4). (C) Heatmap showing tyrosine kinase – substrate relationships. Correlation of phosphorylation site behavior was performed using SELPHI on the combined phosphoproteomics experiment data. Soluble tyrosine kinases (known kinase activating phosphorylation sites are marked with a star) are listed at the top and potential substrates on the left. Positive correlation is indicated in blue while negative correlations are shown in red.

This global signaling analysis identified 33,296 unique phosphorylated peptides from 5,518 proteins corresponding to 27,143 individual phosphorylation sites (Table S1). Significant abundance change (p-value<0.001, Cox and Mann 2008), was observed for 6091 peptides (from 2,314 proteins) in at least one dataset. Since, there is a lack of prior information, with respect to the Fes kinase’s function, substrates and interactome, we used a data driven computational approach to place it in a functional context. Specifically, all acquired quantitative ratio data from the 13 large-scale experiments was pooled and analyzed using SELPHI (Petsalaki et al., 2015) to uncover kinase to substrate dependencies and associations. The SELPHI pipeline assigns a potential kinase-substrate connection based upon a correlation of the level of kinase phosphorylation to the level of phosphorylation on potential substrates assuming a linear relationship. 693 associations were calculated using SELPHI from the 1742 unique tyrosine phosphorylation sites identified by MaxQuant. The predicted connections between activating sites on tyrosine kinases and potential substrates are shown in Figure 2C and Table S2. Many assigned substrate sites were associated with multiple kinases. Such redundancies, as seen for example in shared substrate pools for Lck and Tec (Figure 2C), might be important factors that grant signaling networks a high degree of plasticity and adaptability. Additionally, enrichment of serine/threonine phosphorylated peptides enabled us to calculate SELPHI connections between serine/threonine kinases and potential downstream substrate sites (Table S3). Thus, the perturbation of different signaling components using kinase inhibitors and exogenous Fes kinase expression allowed us to predict global kinase-substrate relationships and extract a potential substrate catalogue for the Fes kinase for subsequent verification.

Twelve potential substrate sites showed a high correlation to Fes activation. A kinase assay was developed using a 13-mer peptide spot array of 100 potential tyrosine sites that included those 12 potential sites and an additional 88 sites predicted from the SELPHI analysis to be linked to activity-driving sites on other soluble tyrosine kinases. The known Fes substrate Cttn as well as the Fes auto-phosphorylation site were included as positive controls (Figure 3A, Table S4). The assays were carried out using both soluble SH2-kinase and isolated kinase domain to test for effects of the SH2-domain on substrate specificity. Similar phosphorylation patterns and a Fes substrate consensus sequence were observed for both constructs (Figure 3A) with 10 of the 12 SELPHI-predicted Fes substrate sites phosphorylated *in-vitro* (Figure 3B). This analysis showed that Fes can readily phosphorylate sites in the activation loop of the tyrosine kinases Syk (Y352), Lyn (Y397), Blk (Y389) and Tec (Y519), which is critical for their activation. Mapk1 Y187 and Stam2 Y374 were not confirmed as Fes substrates (Figure 3B). In addition to Fes and its 4 putative tyrosine kinase substrates identified above, 5 other soluble tyrosine kinases (Fgr, Lck, Abl2, Btk and Tnk2) displayed increased phosphorylation after BCR cross-linking (Figure 3C). From this initial level of tyrosine signaling, the derived network activated the downstream serine/threonine MAP kinases such as Mapk1 (pY187) and Mapk3 (pY204). Our data suggests that Fes does not phosphorylate these components directly, but instead influences mitogenic signaling indirectly through activation of Blk, Syk, Lyn and Tec (Figure 3D). The SELPHI correlation analysis of the data predicts further connections to serine/threonine kinases such as Chek1 and Gsk3b and importantly includes previously established kinase connections such as Mapk1 to Rps6ka1 (Alexa et al., 2015), providing further credence that this phosphoproteomics strategy successfully assembled a directional signaling network. Connecting Fes activity to this extensive network context directed further detailed and mechanistic studies of its influence on B-cell regulation.

**Figure 3.**
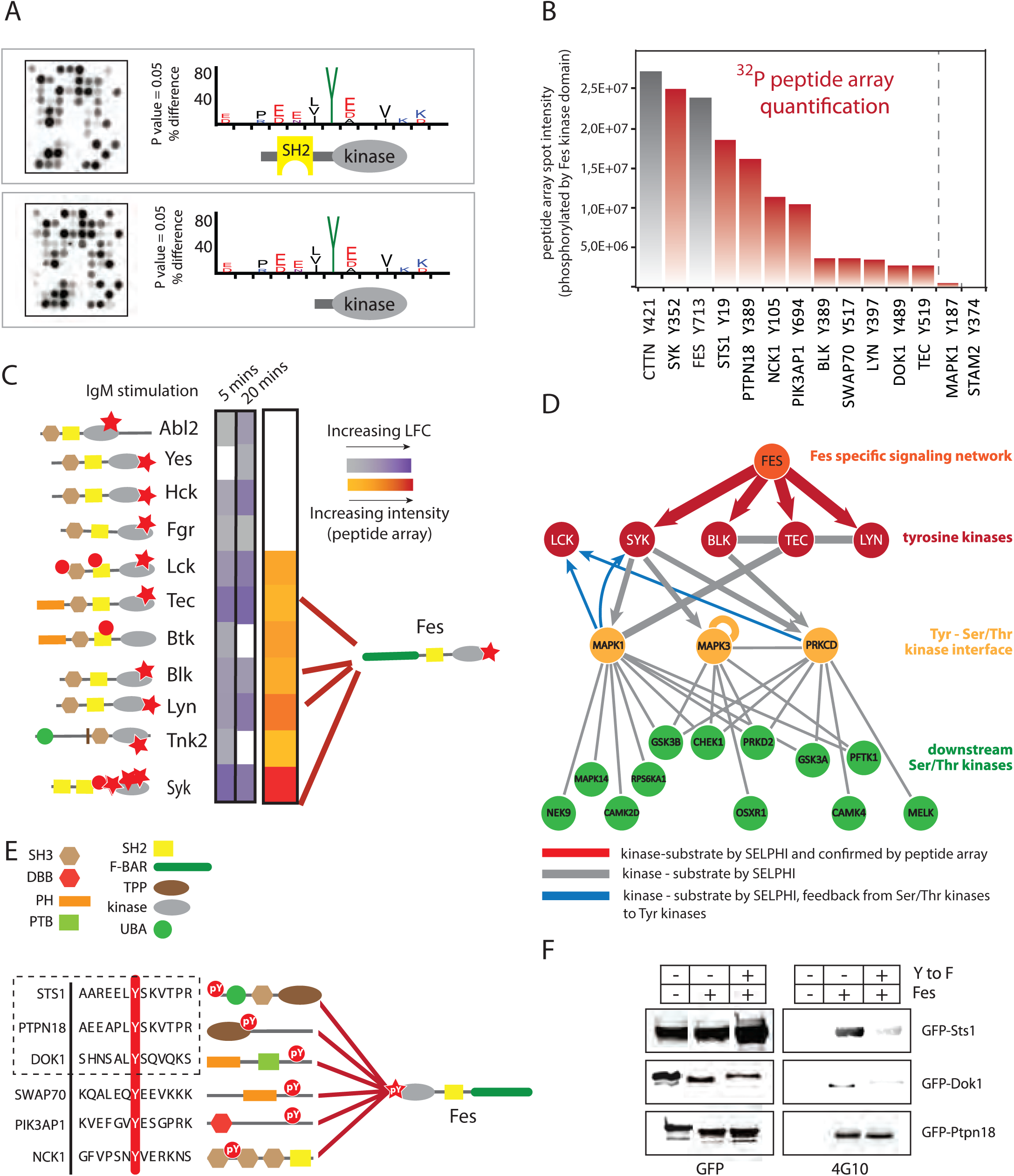
Fes substrates and influence on B-cell kinase signaling networks. (A) Representative examples for radio kinase assays conducted on spots blots of synthesized peptides corresponding to 100 sites exhibiting correlations with activating sites on soluble tyrosine kinases by the SELPHI analysis. Phosphorylation profiles for purified Fes SH2-kinase and kinase domain alone were obtained using Icelogo (Colaert et al., 2009) p-value indicates significance of enrichment of motif compared to all peptides in the array (Table S4). (B) Amount of tyrosine phosphorylation in our peptide array of 12 SELPHI-identified Fes substrate sites compared to known Fes phosphorylation sites (positive controls, CTTN Y421 and Fes Y713 autophosphorylation site, are indicated in grey). Sites that did not meet the phosphorylation intensity cutoff are indicated by the dashed line. (C) Tyrosine kinases that were found to be activated following BCR stimulation. Stars indicate activating tyrosine phosphorylation sites. Red connections symbolize kinase–substrate connections of the Fes kinase that were identified by SELPHI and confirmed via the peptide array. For visual simplicity, gray/purple columns indicate maximum log fold change (base 2) for conditions that represent Fes activation after 5 mins (m/l 5, 6, 8, h/l 8, h/m 12 and h/m 13 in Table S1) and 20 mins IgM (m/l 1, 2, 3, 4 in Table S1) stimulation and orange/red column indicates value from peptide array, where the experiment was performed. White values indicate missing data. (D) Schematic showing the flow of kinase signaling based on our SELPHI analysis, indicating that Fes is capable of influencing kinase components of the phosphotyrosine signaling network but is not directly manipulating components of the Tyrosine kinase – Serine/Threonine kinase interface or downstream kinase network (S/T kinase –Table S3). Arrows are included where there is confidence of directionality of predicted association (E) Non-kinase proteins that were found to be Fes substrates are depicted with the particular phosphorylation site indicated. Sts1, Ptpn18 and Dok1 are circled as they were used for further IP-MS experiments. (F) *In-vitro* phosphorylation of Dok1, Sts1 and Ptpn18 (and their Y-to-F substituted counterparts) by purified Fes kinase. GFP-tagged proteins were expressed in DG75 cells, immunoprecipitated and phosphorylated using purified Fes SH2-kinase for 10 minutes. Kinase reaction was analyzed by immunoblotting (4G10) to assess the phosphorylation state.

### Fes phosphorylation recruits the tyrosine kinase Csk into distinct signaling complexes

In addition to the four tyrosine kinases, we confirmed Fes substrate sites in the guanine exchange factor Swap70 (pY517), various adaptor proteins (Dok1 pY489, Pik3ap1 pY694 and Nck1 pY105) and tyrosine phosphatases (Sts1 pY19 and Ptpn18 pY389) (Figure 3B). The sites on Dok1, Sts1 and Ptpn18 shared a matching phosphorylation motif (LYSxV) (Figure 3E) that prompted further experiments. To confirm direct Fes-mediated phosphorylation of these specific sites, we expressed GFP-tagged WT Dok1, Ptpn18 and Sts1 or tyrosine to phenylalanine mutants of this motif in HEK293T cells, and following anti-GFP antibody immunoprecipitations, performed *in-vitro* phosphorylation assays using purified Fes SH2-kinase. All three proteins were Fes substrates as detected by Western blotting using anti-phosphotyrosine antibody 4G10 (Figure 3F). WT and Y389F mutant Ptpn18 exhibited similar anti-phosphotyrosine signals indicating multiple sites are likely phosphorylated *in-vitro*.

To gain insight into the importance of the LYSxV motif within Dok1, Sts1 and Ptpn18, we expressed both WT and the Y/F mutants in DG75 cells together with the activated Fes E708A kinase mutant to increase phosphorylation on the target proteins and monitored changes to the BCR signaling network by IP/MS (Figure 4A and Table S5). This analysis revealed that the Src-inhibiting tyrosine kinase Csk was recruited to this motif in a Fes-dependent manner as it was only seen in immunoprecipitates with WT Dok1, Sts1, and Ptpn18. Co-expression of the Fes E708A variant led to a constitutive association between all three bait proteins and Csk, with an increase in the presence of IgM stimulation (Table S5), which could be abolished by the Y/F mutations.

**Figure 4.**
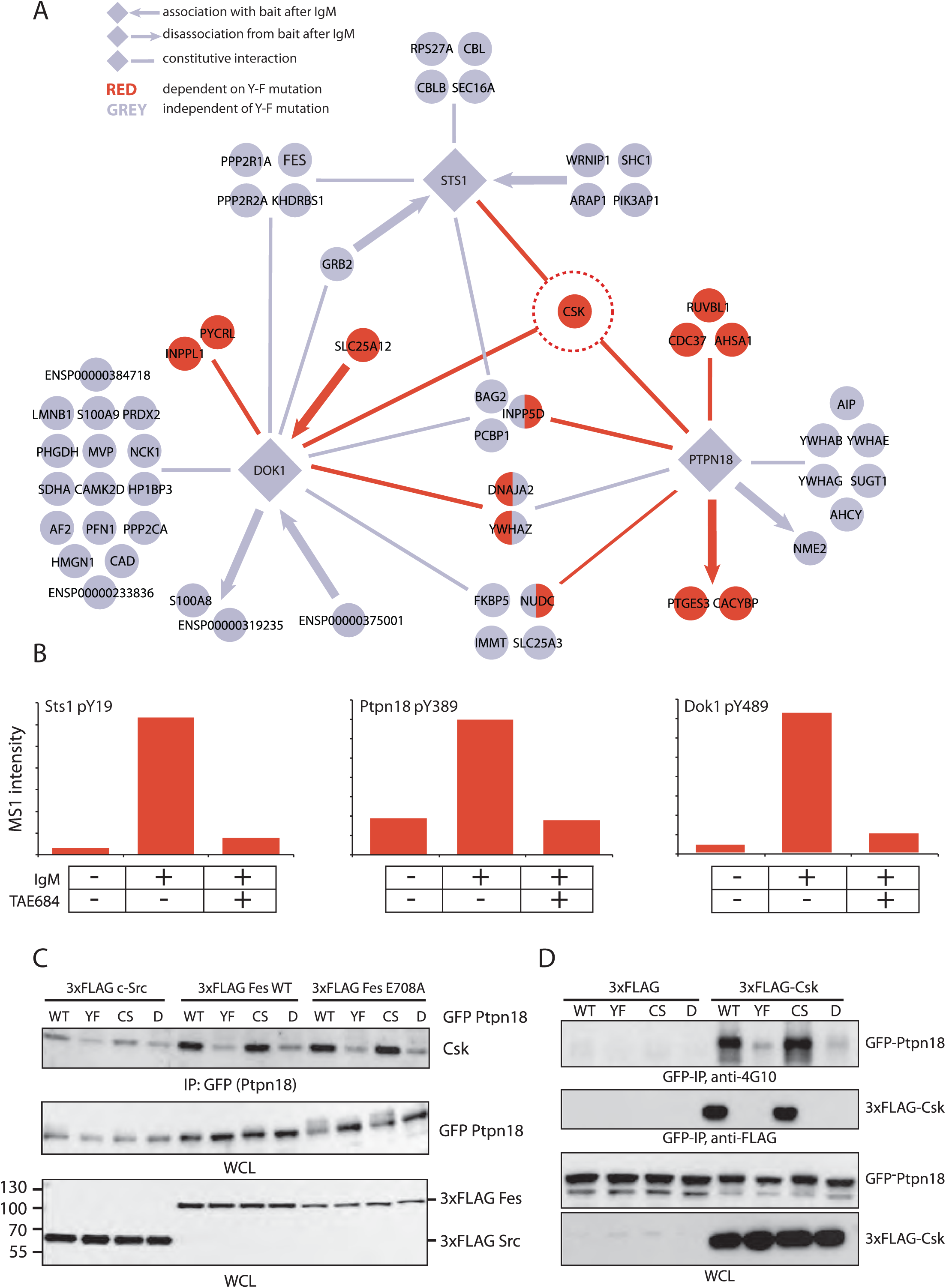
Dynamic interactome analysis reveals Fes-mediated Csk recruitment. (A) Y-to-F substitutions of Fes phosphorylation sites were employed to study the interactome of Dok1, Sts1 and Ptpn18, revealing Y-F (Fes phosphorylation) dependent (red) as well as Y-F independent and constitutive interactors (grey). Arrows towards the bait indicate association and arrows away from the bait disassociation following 10 minute BCR stimulation. Increase/decrease upon stimulation has not been mapped on the figure for simplicity. Identified peptides for the respective baits are reported in Table S5. Ion intensities of endogenous Sts1 pY19, Ptpn18 pY338 and Dok1 pY449 tyrosine-phosphorylated peptides in resting DG75 cells and following BCR cross-linking using anti-human IgM. The peptides were labeled using dimethyl labeling and Fes phosphorylation was inhibited using a pre-treatment with the TAE684 compound. (C) Co-expression of FLAG-tagged tyrosine kinases (Src, Fes, Fes E708A) with GFP-Ptpn18 showed that Fes but not Src is capable of causing Csk-Ptpn18 association. Mutation of Y388 to F (YF) on Ptpn18 abolished the interaction while the mutation of C228 to S (CS, phosphatase catalytic residue) did not affect Csk recruitment, double mutation indicated as D. (D) Co-transfection of Flag-Csk with GFP-Ptpn18 in Hek293T cells showed that expression of Csk leads to association with and phosphorylation of Ptpn18 which is dependent on Ptpn18 Y388.

In our large-scale phosphoproteomics experiments, the Fes-specific inhibitor TAE684 effectively blocked phosphorylation on these sites during BCR stimulation experiments in WT DG75 cells (Figure 4B). To further demonstrate kinase specificity at these sites, we tested co-expression of Ptpn18 with c-Src and c-Fes in Hek293T cells and found that the latter kinase orchestrated a more robust interaction between Csk and Ptpn18, consistent with direct substrate connection of Fes with Ptpn18 (Figure 4C). Co-expression experiments of Csk and Ptpn18 showed that Csk is also able to directly or indirectly induce tyrosine phosphorylation of this binding partner and that this phosphorylation is dependent on the Fes target site (Figure 4D), emphasizing the Fes phosphorylation site as an initiation point for further signaling events.

### Fes signaling impacts CD19 levels via its substrate Sts1

Following the observation that the Fes kinase co-localizes to the BCR and can be activated following BCR stimulation, we performed a STRING (Szklarczyk et al., 2019) protein network analysis on BCR components and found that the interactome of Fes substrate Sts1 is tied to the regulation of BCR receptor proteins (CD79A, CD79B and CD19) via the E3 ubiquitin ligase Cbl (Figure 5A). CD19 plays a decisive role in B-cell activation as it lowers the threshold for the initiation of the signaling response upon exposure to an antigen. A successful receptor engagement will lead to CD19 phosphorylation on cytoplasmic tyrosine residues such as Y531 which in turn serve as docking sites for the regulatory subunit p85 of PI3K (Otero et al., 2001), (Depoil et al., 2008). We observed that titration of WT Fes kinase into DG75 cells resulted in a complex alteration of endogenous CD19 protein levels with a distinct increase at low exogenous kinases levels and reduction at higher Fes expression (Figure 5B). Csk-mediated Src inhibition may be involved in this modulation of CD19 since phosphorylation of the inhibitory Src Y527 site, a known substrate for the Csk tyrosine kinase, paralleled the observed changes in CD19 levels. This effect was Fes-specific as overexpression of c-Src or Tec did not produce this effect on pY527 and CD19 protein levels (Figure 5B).

**Figure 5.**
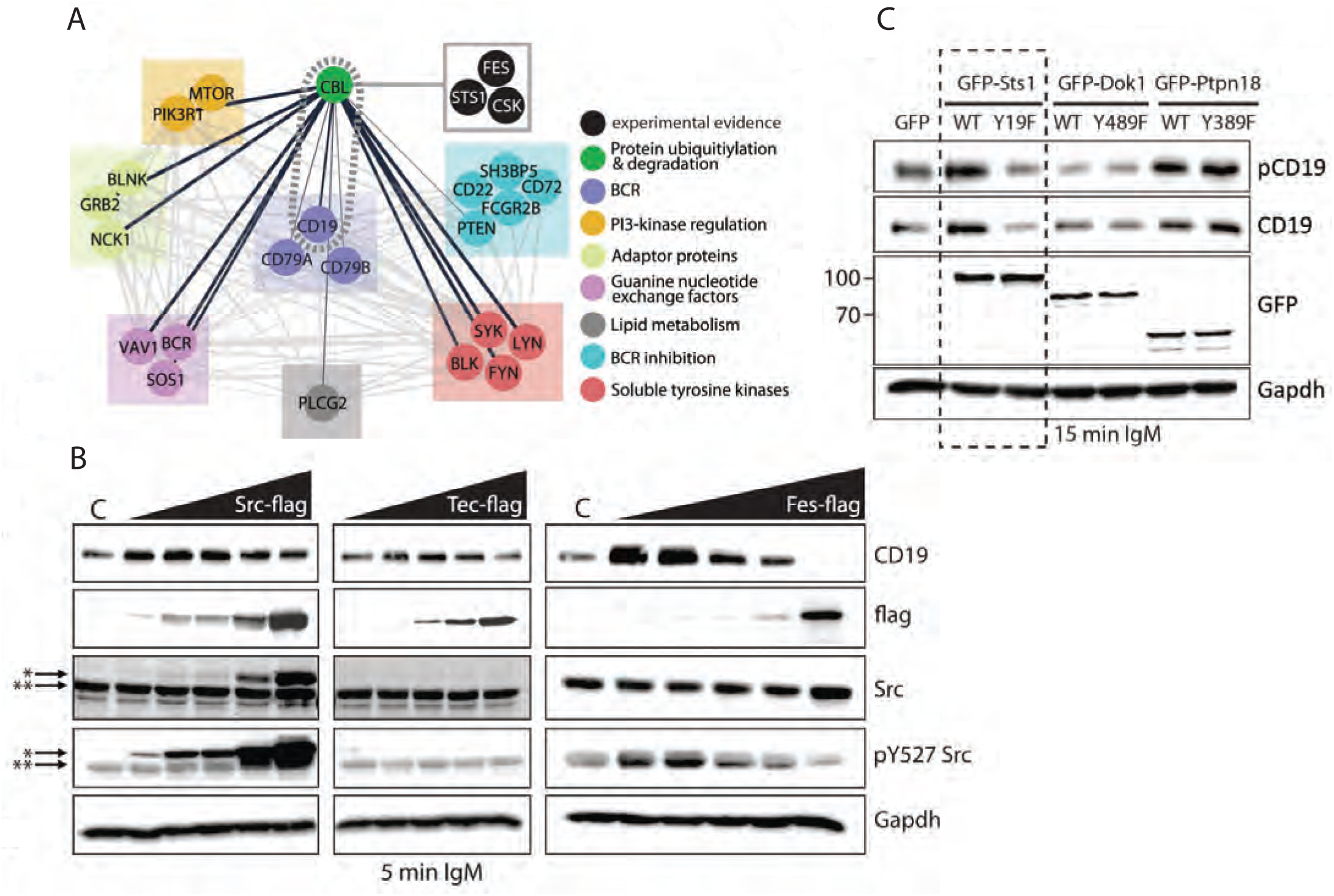
Fes modulates CD19 protein levels via its substrate Sts1. (A) STRING (http://string-db.org/) protein association analysis performed on BCR components CD79A/B and CD19 revealed a connection to Fes/Sts1/Csk directed signaling pathways via the E3 ubiquitin ligase Cbl. (B) Transient expression of Flag-tagged E708A Fes, Tec and Src in BCR-stimulated DG75 cells. Kinase expression levels were titrated (1, 5, 20, 50,100 ug of plasmid DNA) while keeping the total amount of transfected DNA constant. * marks the exogenous Src protein while ** indicates endogenous Src Protein levels, as indicated, as well as the Csk-mediated Src phosphorylation site pY527 were analyzed by immunoblotting. (C) GFP-tagged Dok1, Sts1 and Ptpn18 and their counterparts with mutations of the Fes phosphorylation sites were expressed in DG75 cells and stimulated with IgM. Effects on CD19 levels were analyzed by immunoblotting. Only the Y to F mutation on Sts1 led to a change in CD19 protein levels (indicated by the dashed line).

Since we identified Sts1, Dok1 and Ptp18 as Fes substrate proteins that mediate Csk recruitment, we also tested their ability to modulate CD19 levels. We ectopically expressed WT versions of these proteins or derivatives where the identified Fes tyrosine phosphorylation sites were inactivated through mutation to phenylalanine and monitored CD19 levels (Figure 5C). Only Sts1 displayed a difference related to the Fes substrate site as the WT protein generated higher CD19 levels than the mutant did. To monitor for an effect on CD19 function in coordinating downstream PI3 kinase and Src signaling pathways, we also measured phosphorylation on Tyr531 of CD19 using a phospho-specific antibody (Figure 5C). While differences in signal were observed, they paralleled those seen for CD19 levels. This indicates that the Fes-mediated Sts1 phosphorylation modulates the magnitude of the B-cell response by altering the overall level of the CD19 protein.

## Discussion

Dysregulation of kinase function accounts for a large number of pathologies from cancer to immune diseases (Lahiri et al., 2010). Depending on the cancer type, several kinases may act either as an oncogene or a tumor suppressor (An & Brognard, 2019), demonstrating the intricacies of kinase regulation and their role in defining cell phenotypes. Despite kinases being excellent targets for drug intervention (Bhullar et al., 2018), the majority of them, including the Fes kinase, are understudied (Invergo & Beltrao, 2018), thus making it difficult to link their function with cell phenotype. Here we have applied an unbiased phosphoproteomic and data-driven strategy to shed light on the contrasting oncogenic and tumor suppressor function of the understudied Fes kinase.

### Fes signaling involves localization and SH2 / kinase-domain cooperativity

Antigen binding to the BCR induces clustering of the receptor into lipid rafts (Gupta and DeFranco, 2007) and facilitates the downstream signaling cascade. The Fes F-BAR domain seems to be the driving force for Fes membrane localization (McPherson et al., 2009) and we demonstrate that it also affects substrate selectivity. The decrease in Fes activation, when the F-BAR domain is deleted, points to membrane localization and dimer-/oligomerization being a critical factor for signal transduction to Fes after BCR clustering. Hypothetically, membrane rearrangement during BCR clustering could concentrate BCR-associated Fes kinase into signaling environments leading to its auto-phosphorylation and activation. This then in turn, would lead to phosphorylation of substrates such as membrane-associated tyrosine kinases (e.g. Syk, Lyn), phosphatases (Ptpn18, Sts1) and adaptor proteins such as Dok1, that further direct cellular fate.

The SH2 domain was originally defined in Fes as a non-catalytic region contributing to the catalytic activity of the neighboring kinase domain (Sadowski et al., 1986) with whom it forms a stable complex (Koch et al., 1989) (Weinmaster and Pawson, 1982). Interestingly, our identified consensus sequence for Fes kinase specificity matches the Fes SH2-binding specificity (YEXVX) as determined previously (Filippakopoulos et al., 2008). The SH2 domain links phosphotyrosine binding of primed targets to catalytic activation (Filippakopoulos et al., 2008). However, it is also conceivable that shared motif specificity of the SH2 and kinase domain for the same residue leads to distinct signal amplification by increasing kinase concentration in a particular cellular microenvironment with highly abundant substrates through a synergistic effect of phosphorylation and localization. In this regard, it has been demonstrated that Dok1 and Sts1 can form oligomers in cells (Kowanetz et al., 2004), (Songyang et al., 2001) underlining the possibility of signal amplification through overlapping of kinase and SH2 specificity.

We demonstrated that Fes phosphorylation in B-lymphocytes of Dok1, Sts1, and Ptpn18 recruits the Csk tyrosine kinase. Csk is not primarily regulated by phosphorylation; instead it seems to be dominantly spatially controlled (Vang et al., 2003) by phosphorylation-based recruiting via its SH2 domain. Our data indicates that the Csk kinase and Fes kinase SH2 specificity have evolved to complement each other by amplifying phosphorylation at the very residues they are supposed to bind (Figure 4A/4D). In-vitro determination of SH2-domain specificity also found similar consensus motifs for Csk and Fes SH2-domain binding (Huang et al., 2008). This highlights again the close association between those two protein domains in the regulation of signaling circuits.

Taken together, our results point to a model where Csk localization is initiated by Fes phosphorylation on specific protein complexes and amplified by the SH2 and kinase domains of both kinases. Consecutively, this would lead to a spatially and temporally controlled, localized Src inactivation within specific protein complexes in the cell.

### The Fes kinase – tumor suppressor or oncogene?

Although our SELPHI analysis showed that Fes is able to activate downstream mitogenic kinases such as Mapk1/3 through signal propagation, our current evidence points to Fes’s involvement in the modulation of the critical decision process around Src activation and inhibition as a possible explanation of its seemingly paradox functions in cancer. We show that Fes signaling plays a crucial role in tuning Src kinase activity in a cellular model system: titration of Fes expression levels indicated that Fes concentration is critical to tip the scales of Src regulation via Csk recruitment in either direction, which may contribute to the resulting modulation in CD19 levels that are concomitantly observed (Figure 5A).

Sts1, a ubiquitin-binding tyrosine phosphatase, has been implicated in counteracting Cbl-mediated receptor degradation through the dephosphorylation of target sites for the Cbl SH2 domain that guide the receptor ubiquitination and degradation process (Kowanetz et al., 2004) (Raguz et al., 2007). Typically, these sites are Src targets (Mikhailik et al., 2007) and it is conceivable that Csk recruitment to Sts1 signaling complexes is needed to locally inhibit Src activity in cooperation with the Sts1 dephosphorylation of Src target proteins to facilitate its function. Additionally, it has been demonstrated that CD19 can form a complex with Cbl, down-regulating activated CD19-PI3K complexes (Beckwith et al., 1996). We found that Csk localization to Sts1 via Fes phosphorylation is necessary for Sts1 function of stabilizing CD19, possibly by opposing Cbl function.

Previous data on enforced CD19 expression showed that, despite its function to balance the B-lymphocyte signaling threshold, CD19 also leads to growth retardation of malignant plasma cells (Multiple myeloma) (Mahmoud et al., 1999) and thus can act as a tumor suppressor in the lymphoid system. As such, it is an important factor in lymphoid malignancies since CD19 loss is often observed in Multiple myeloma cancerous cells (Mateo et al., 2008), (Cannizzo et al., 2012). Our data indicates that low Fes expression levels in lymphocytes might constitute a signaling threshold where Fes-mediated Csk recruitment contributes to high CD19 levels, helping to safeguard lymphocytes from tumorigenic degeneration. Interestingly, rising Fes kinase ectopic expression levels can also have an opposite compensation effect, strongly reducing CD19 levels. This observation might provide a novel angle for therapeutic strategies targeting Fes using kinase inhibitors to reverse such repressive effects on CD19 in plasma cell cancers. In support of this, an RNAi screen demonstrated Fes to be an important driver of multiple myeloma cell survival (Tiedemann et al., 2010).

By lowering CD19 levels, Fes might contribute to an oncogenic phenotype when expressed at high levels in plasma cells but its ability to control Csk recruitment could be a means to counteract oncogenic Src signaling in other tissues. This might be especially pronounced in colon cancer where aberrant Src signaling has been detected in about 80% of the cases (Chen et al., 2014) and where Fes has been identified as a tumor suppressor. Additionally, it has been shown that Src can prime the adaptor Shc1 for EGFR phosphorylation thus activating the Ras-MAPK pathway (Begley et al., 2015). In such a molecular context, it is conceivable that Fes expression might indeed help to counteract oncogenic Src and thus the Fes kinase would operate as a tumor suppressor.

In summary, we present a systems biology approach to shed light on the enigmatic biological role of the Fes kinase. We report that Fes is able to intricately modulate Src signaling, recruiting the Src inhibitory kinase Csk to multiple signaling complexes in lymphocytes. We observed that this mechanism controls the protein levels of the CD19 co-receptor, at least in part, via the Fes substrate Sts1. We propose that, through dysregulation of these mechanisms, Fes could potentially contribute to oncogenic effects in the lymphoid system and serve as possible novel drug target in specific cancers such as CD19-deficient multiple myeloma.

## Methods

### Cell culture, transformation and stimulation

The DG75 human lymphoblast cell line and the mouse macrophage RAW264.7 cell line were obtained from American Type Culture Collection (ATCC). DG75 cells were cultured at 37°C, 5% CO_2_ in RPMI with glutamate/Pen-Strep supplemented with 10% fetal bovine serum (FBS). For transfection, cells were grown to a density of 0.7×10^6^ and washed 3 times in RPMI. 20ug of plasmid DNA was mixed with 10^7^ cells in 0.5ml of RPMI. Cells were incubated for 10 minutes and electroporated (BTX Electro Square Porator ECM 830) by three 8ms pulses at 225V in a 4mm cuvette. After 10 minutes of incubation, cells were placed into RPMI with 10% FBS. For transient transfections, cells were then grown for 48 hours. For the generation of stable cell lines, Geneticin (G418, Life Technologies) was added at a final concentration of 1mg/ml after 48 hours. After 2 weeks of culturing, cells stably transfected with Fes-GFP constructs were FACS sorted and the highly GFP-labeled cell population was expanded and maintained in Geneticin containing RPMI medium. RAW264.7 cells were cultured in Dulbecco’s Modified Eagle’s Medium supplemented with 10% FBS.

For BCR stimulation experiments, DG75 cells were starved using RPMI medium without FBS for 2 hours. For kinase inhibition, cells were incubated for 1 hour with respective kinase inhibitor (1uM final concentration). Cells were pelleted and resuspended in 1ml of RPMI. Goat anti-human IgM was added at an amount of 1 microliter of a 10 mg/ml solution per 1 million cells. After incubation at 37°C, cells were put on ice, pelleted and frozen on dry ice prior to analysis.

### Gateway cloning and Openfreezer repository

A detailed cloning summary is provided in Table S7 including acknowledgements of clones from external sources. To create gateway entry clones, Open Reading Frames (ORFs) were amplified by polymerase chain reaction (PCR) from the templates indicated in Table S7 using Phusion DNA polymerase (NEB) with Gateway compatible sequences appended to the end of the primers (5’ sequence - gggg aca act ttg tac aaa aaa gtt ggc acc, 3’ sequence - gggg ac aac ttt gta caa gaa agt tgg gta). The resulting products were then cloned into pDONR223 (Rual et al., 2004) using BP clonase (Invitrogen) and subsequently into Gateway destination vectors using LR Clonase (Invitrogen) according to the manufacturer’s protocols. All inserts were sequence verified using CodonCode Aligner software. Clones were tracked in OpenFreezer (Olhovsky et al., 2011). Point mutations were generated in relevant expression vectors using the QuikChange methodology according to the manufacturer’s instructions (Agilent Technologies).

### Immunofluorescence microscopy

DG75 cells were starved for 1 hour in RPMI without FBS causing adherence of the cells to glass cover slips. Cells were fixed with 2% paraformaldehyde in PBS for 10 minutes. Cells were washed 3 x 5 minutes with PBS. Permeabilization was performed by incubation with 0.5% Triton-X100 in PBS for 5 minutes. Cells were washed 3 x 5 minutes in PBS. Blocking was performed for 1 hour with 5% BSA and 0.1% Tween-100 in PBS. Primary antibody labeling was performed in 1% BSA using an anti-Fes antibody (Cell Signaling Technology) and goat anti-human IgM to label the B-cell receptor. Cells were washed 3 x 5 minutes in PBS. Secondary labeling was performed using a donkey anti-rabbit Alexa Fluor 647 (Abcam) and a donkey anti-goat Alexa Fluor 488 (Abcam) in 1% PBS for 1 hour. The actin cytoskeleton was labeled simultaneously with Alexa Fluor 555 Phalloidin (Life Technologies). Cells were washed 3 x 5 minutes in PBS and cover slips were rinsed with water and mounted face down on glass slides. Samples were then imaged on an Inverted Leica DMIRE-2 microscope equipped with fluorescence optics. Using either a 60X- or 100X-oil immersion lens, fluorescent images were projected onto a Hamamatsu ORCA CCD camera and captured using VOLOCITY software (Improvision).

### FSBA assisted kinase assays

5’-(4-Fluorosulfonylbenzoyl) adenosine hydrochloride (FSBA) was obtained from Sigma-Aldrich. 1×10^7^ DG75 cells were pelleted and lysed in NP40 buffer containing cOmplete, EDTA free protease inhibitor cocktail (Roche) by passing them through a 25-gauge needle and three pulses of sonication. Cell debris was removed by a 10000g centrifugation step for 10 minutes. Protein concentration was adjusted to 5mg/ml. FSBA was dissolved in DMSO (25mg FSBA in 500ul DMSO) and added to the cell lysate in a 1(FSBA):5 (lysate) ratio. The mixture was incubated at 30°C for 1 hour. DMSO was diluted to 5% with NP40 buffer and precipitated FSBA was removed by centrifugation for 2 minutes at 10000g. Sample volume was concentrated in YM-10 spin columns and adjusted to 3mg/ml. For the kinase assay, the kinase inhibited lysate was diluted 1:1 with 40mM MOPS, pH 7.2, 50mM glycerophosphate, 10mM EGTA, 2mM NaVO_4_, 2mM DTT, 50mM MgCl_2_, 2mM ATP. Purified kinase (full length Fes and SH2-kinase) was added (equal amount of activity) and assayed at 30°C for 30 min. 20ug of protein lysate was subsequently analyzed by blotting with anti-phosphotyrosine antibody (4G10).

### Multiple reaction monitoring (MRM) mass spectrometry

Raw data files from data-dependent acquisition (DDA) experiments were imported into MRMpilot 1.1 (SCIEX) software to generate a list of potential sMRM transitions. Peptide retention times were determined by MS/MS from multiple MRM Initiated Detection and Sequencing (MIDAS) analyses using MultiQuant (version 2.1) software. The final sMRM method consisted of 54 transitions from the Fes kinase. For data normalization, we chose 5 Fes peptides that were unlikely to undergo post-translational modifications. sMRM analysis was performed on a hybrid triple quadrupole/ion trap mass spectrometer (5500 QTrap; SCIEX). Chromatographic separation of peptides was carried out on a nano-LC system (Eksigent, Dublin CA) coupled to a 100 μm i.d. fused silica column packed with 5μm ReproSil-Pur C18-AQ as a trap column and a 75 μm i.d. fused silica column packed with 3 μm Reprosil-Pur C18-AQ as the separation column (pulled to ∼ 10 μM) to act as an electrospray emitter. Peptides were separated with a linear gradient from 2-30% acetonitrile in 90 min at a flow rate of 300 nl/min. The MIDAS workflow was employed for sMRM transition confirmation. Each sMRM run was scheduled using previously determined LC retention times with a 5-minute MRM detection window and a 3-second scan time with both Q1 and Q3 settings at unit resolution. The MS/MS spectra acquired by MIDAS were first searched against relevant Ensembl databases using Mascot to confirm the identities of peptides. Then the raw data was imported into MultiQuant v2.1 (SCIEX) for automatic MRM transition detection followed by manual inspection by the investigators to increase confidence. Subsequently, the extracted Ion Chromatogram (XIC; proportional to peptide abundance) of each transition was calculated.

### Cell lysis and dimethyl labeling

DG75 cell pellets were frozen on dry ice. Cells were lysed in 50mM ammonium bicarbonate, 8M urea, EDTA free cOmplete protease inhibitor cocktail (Roche), 2mM vanadate, phosphatase inhibitor cocktail 3 (Sigma-Aldrich). Three 20s pulses of sonication were used to solubilise the material. Proteins were reduced by the addition of dithiothreitol (final concentration of 20mM) for 30 min at room temperature. Alkylation was performed by adding iodoacetamide (50mM final concentration) and incubation for 30 min. Urea concentration was subsequently reduced to 2M and proteins were digested overnight at 37°C after the addition of trypsin (Sigma-Aldrich) at a 1:100 ratio. A second round of digestion was performed for 2 hours. Samples were acidified with formic acid (5% final volume) and forming precipitate was removed by centrifugation. Peptides were bound on C18 Sep Pak cartridges (Waters) and desalted with 5% formic acid. Chemical dimethyl labeling was performed on column (Boersema et al., 2009). Samples were combined after elution and freeze dried.

### Phosphopeptide analysis

90mg of dried combined peptide sample was split into two equal batches. With the first batch, immunoprecipitation of tyrosine phosphorylated peptides was carried out. Peptides were dissolved in 5ml IP buffer (50mM Tris, pH 7.4, 150mM NaCl, 0.6% NP-40). 80ul of anti-phosphotyrosine (4G10) agarose beads (Millipore) were added and incubated for 2 hours. Beads were subsequently washed three times with IP buffer and 3 times with water. Peptides were eluted two sequential times with 0.2% TFA and the eluates were combined and freeze dried before mass spectrometry analysis.

The second batch was dissolved in 0.3% TFA, 25% lactic acid, 60% acetonitrile. Titansphere TiO_2_ material (GL Science) was added in a peptide to TiO2 ratio of 1:6 and phosphopeptides were enriched according to the Titansphere manual instructions. After sequential elutions with 1% ammonia and 1% pyrrolidine, eluates were combined and the solvent evaporated. A desalting step using Sep Pak columns (Waters) was performed before the phosphopeptides were freeze dried and reconstituted in 30% acetonitrile, 5mM KH_2_PO_4_, pH 2.7. Peptide fractionation was carried out using a PolySULFOETHYL A column (PolyLC inc.), (Gauci et al., 2009). Two minute fractions were collected until 60 min and subsequently freeze dried. Desalting of the fraction was carried out using C18 stage tips (Thermo Scientific) according to the stage tip description. The organic eluent solvent was evaporated and fractions were analyzed on a Orbitrap Velos mass spectrometer coupled to a 1D+ Nano LC (Eksigent). Samples were injected directly onto a column packed in-house (75 μm inner diameter) with 3.5 μm Zorbax C-18 (Agilent) and analyzed using a 90 min gradient from 5%-30% solvent B (Solvent A: 0.1% formic acid in water; Solvent B: 0.1% formic acid in acetonitrile) at a flow rate of 250 nl/min. Raw data was processed using MaxQuant v.1.1.1.36. Mass tolerance for the first search was set to 20ppm. Oxidation (M) and phospho (S/T/Y) were set as variable modifications while Caramidomethyl (C) was set as fixed modification. Peptide and site false discovery rate were set to 0.01. All other settings were kept as default. Data was searched against the human Refseq database v.42 (Pruitt et al., 2014) and the obtained phosphopeptide evidence (Table S1) was subjected to SELPHI analysis. The mass spectrometry proteomics data have been deposited to the ProteomeXchange

Consortium via the PRIDE partner repository (Vizcaino et al., 2016) with the dataset identifier PXD005332 (username; password: w1BLTp6y).

Significant changes in the log2 silac ratios were estimated using the FDR corrected significance A values (Cox and Mann 2008) with a p-value significance cut-off of 0.05, implemented using R software.

### SELPHI analysis

Replicate sample and conditions were merged by keeping the ratio that indicated the maximum level of change in the phospho-peptide intensity. The protein sequences on which the peptide locations were mapped were extracted from the Human RefSeqV42 (Pruitt et al., 2014) and the gene names were mapped using UniprotKB release 2014_09 (UniProt Consortium, 2014). We restricted the dataset to peptides that appear in at least six conditions, i.e. 2 samples and we applied a SELPHI-analysis (Petsalaki et al., 2015) using the default parameters (Spearman correlation cutoff = 0.5 and p-value of correlation 0.05) on the combined datasets. SELPHI identified pairs of kinases associated with the identified phospho-peptides (Tables S2/3).

### Fes kinase purification

cDNAs encoding human Fes (NP_001996) were cloned into pNIC28-Bsa4. Expression constructs were transformed into phage-resistant *E. coli* BL21(DE3)-R3 co-transformed with an expression vector encoding *Yersinia* phosphatase YopH ; :. Cells were grown and Fes SH2-kinase proteins were purified according to previously described procedures (Filippakopoulos et al., 2008).

### Peptide array and radioactive kinase assays

Peptide arrays were prepared using an automated Multipep synthesizer (Intavis) on a cellulose support with standard F-moc chemistry protocols as recommended by the manufacturer and previously described (Warner et al., 2008). Prior to the kinase assay, the membranes were conditioned through a first rinse in 95% EtOH followed by the washing steps, 3x 5 min with TBS-T (20 mM Tris HCl, pH 7.5, 150 mM NaCl, 0.05% Tween 20), 2x 5 min with TBS (20 mM Tris HCl, pH 7.5, 150 mM NaCl) and 2x 5 min with kinase reaction buffer (KRB – 250 mM NaCl, 20 mM MgCl2, 20 mM MnCl2 in 50 mM HEPES buffer pH 7.5). For the kinase reaction, 50 μg of purified Fes kinase or SH2-kinase were used in 5 mL of KRB containing 50 μCi of radio-labeled γ32P-ATP and 40 μM unlabelled ATP. The reaction was stopped by three 20 min washes in Wash Buffer A (8M urea, 1% SDS, 0.5% beta-mercaptoethanol) followed by one 20 min wash in Wash Buffer B (50% EtOH, 10% acetic acid). The membrane was rinsed in 95% EtOH and allowed to air dry prior to exposing XAR film (Kodak) (Warner et al., 2008).

### Immunoprecipitation and MS analysis of protein complexes

DG75 cells were transiently transfected with Gateway expression vectors (Invitrogen) containing GFP tagged Dok1, Sts1 and Ptpn18 constructs. At the same time a flag-tagged Fes E708A gain-of-function construct was also transfected. 48 h after transfection, cells were harvested, stimulated or not for 10 minutes with goat anti-human IgM and lysed in NP-40 buffer. Protein complexes were immunoprecipitated using GFP-Trap (Chromotek) and washed 3 times with 50mM ammonium bicarbonate after the precipitation to remove detergent. Samples were processed using a solid-phase digest protocol as previously described (Bisson et al., 2011). Briefly, columns were made with 200 μm (internal diameter) fused-silica tubing and packed with 3 cm of PolySulphoethyl A beads (particle size 12 μm, pore size 300 Å) (Western Analytical, CA, USA) using a pressure bomb (nitrogen gas, 100-500 psi). Protein eluates were loaded on the column and washed with 10 mM potassium phosphate buffer pH 3 followed by HPLC grade water. Bound proteins were reduced in DTT solution (100 mM DTT/10 mM NH_4_HCO_3_, pH 8) for 30 minutes and washed with HPLC grade water. Reduced proteins were alkylated and trypsin-digested for 1-2 hours in trypsin solution (2 mg/ml sequencing grade porcine trypsin (Promega)/100 mM Tris/10 mM iodoacetamide, pH 8). Peptides were eluted in 15 μl of 200 μM NH_4_HCO_3_, pH 8 and 1 μl of 50% formic acid was added for a final volume of 16 μl. Samples were analyzed on a TripleTOF 5600 mass spectrometer equipped with a nanospray III ion source (AB SCIEX), and coupled to a 1D+ Nano Ultra LC (Eksigent). Samples were injected directly onto a column packed in-house (75 μm inner diameter) with 3.5 μm Zorbax C-18 (Agilent) and analyzed using a 90 min gradient from 5%-30% solvent B (Solvent A: 0.1% formic acid in water; Solvent B: 0.1% formic acid in acetonitrile) at a flow rate of 250 nl/min. Tandem mass spectra were extracted, charge state deconvoluted and deisotoped in Analyst version 2.0. Wiff files were converted to .mgf format using ProteinPilot software and both formats were stored in ProHits (Liu et al., 2010). All MS/MS spectra were analyzed using Mascot (Matrix Science) version 2.2 against the human RefSeq database (release 57 supplemented with entries from the cRAP database from GPM - a total of 72226 sequences; 39792614 residues). Trypsin was chosen for digestion with up to 2 missed cleavages. Carbamidomethyl (C) was set as a fixed modification and oxidation (M), Phospho (ST) and Phospho (Y) were set as a variable modification. ESI-QUAD-Tof type fragmentation was selected with peptide mass tolerance set to 30 ppm, and fragment mass tolerance set to 0.2 Da (peptide identifications in Table S5).

We first extracted the total peptide counts for each identified protein and generated files compatible with SAINT (Choi et al., 2012). The preys were also annotated with the score from the CRAPome database (Mellacheruvu et al., 2013). We only consider as interactors preys with a SAINT score greater than 0.8 (SAINT output can be found in Table S6).

### Csk binding and phosphorylation assays

Mammalian expression vectors encoding Flag-epitope tagged full-length Csk and eGFP-tagged Ptpn18 WT & Y/F mutant were transfected into 293T cells using 5 ug/mL PEI (Polyethylenimine, Sigma Aldrich) in Opti-MEM (GIBCO). Following 18-hour transfection, cells were stimulated with hydrogen peroxide-activated sodium vanadate for 15 min at 37°C. Lysates, prepared in 50 mM Tris HCl pH 7.5, 150 mM NaCl, 3 mM EDTA, 1 % NP40, 10 ug/mL aprotinin, 10 ug/mL leupeptin, 1 mM Na Vanadate, were proportioned for analysis of protein expression (5%) and immunoprecipitation (95%) using GFP-Trap beads (ChromoTek, Germany). Co-immunoprecipitated proteins were detected using anti-Flag antibody (M2 – Sigma), eGFP fusion protein levels monitored using anti-GFP antibody (ab290 – abcam) and phosphotyrosine levels by blotting with anti-phosphotyrosine antibody (4G10) and ECL visualization.

### Western blot

Protein blots were produced by semidry transfer. Visualization of antibody signals was achieved using HRP-conjugated secondary antibodies and the chemiluminescence substrate SuperSignal West Pico (Thermo Fisher Scientific). Blots were imaged and quantitatively analyzed using a Chemidoc MP imager (Biorad). *In-vitro* kinase assays for Sts1, Dok1, Ptpn18 and Cttn were evaluated using an Odyssey Infrared Imaging System (LI-COR) using fluorescently labeled secondary antibodies at 800nm and 680nm. Images were analyzed using the ImageJ software (http://imagej.nih.gov/ij/).

### Inhibitors and antibodies

Src Inhibitor 1, PP2 and PD184352 were obtained from Sigma Aldrich. The TAE684 compound was obtained from the Gray laboratory (Harvard Medical School, Boston).

#### Antibodies against the following proteins were obtained from Cell Signaling Technology

Fes (#2736), Lyn (#2732), phospho-Lyn Y507 (#2731), phospho-Btk Y223 (#5082), phospho-Syk

Y525/526 (#2711), CD19 (#3574), phospho-CD19 Y531 (#3571), Akt (#9272), phospho-Akt S473

(#9271), Erk1/2 (#9102), phospho-Erk1/2 (#9101), GFP (#2555), Src (#2108), phospho-Src Y527 (#2105),

#### Antibodies against the following proteins were obtained from Santa Cruz Biotechnology

Fes (sc-7671), Dok1 (sc-6374), Gapdh (sc-365062)

#### Antibodies obtained from other vendors

Anti-phosphotyrosine 4G10 (Millipore), anti-Flag M2 (Sigma Aldrich), goat anti-human IgM (Southern Biotech), anti-GFP ab290 (Abcam)

### Data availability

The mass spectrometry proteomics data have been deposited to the ProteomeXchange Consortium (http://proteomecentral.proteomexchange.org) via the PRIDE partner repository (Perez-Riverol Y et al, 2019) with the dataset identifier PXD005332.

## Author Contributions

AOH, TP, MK and KL conceived the study. EP and AOH designed the experiments. AOH conducted most experiments and wrote the manuscript with input from EP, KC and GG. MK performed kinase purifications and provided constructs. AOH and LS analysed the raw phosphoproteomics data. EP designed the Proteomics Pipeline and performed most of the bioinformatics analyses and supervised LS. CZ and MT provided technical assistance to mass spectrometric analysis. GG synthesized the peptide arrays and performed Csk binding assays. KC contributed to manuscript preparation and funding. TP provided funding, lab space and facilities.

## Acknowledgements

This work was supported by the Ontario Research Fund Round 5 and GL2 grants to Dr. Tony Pawson. LS and EP are funded by EMBL-EBI.

## Figures

**Figure S1.**
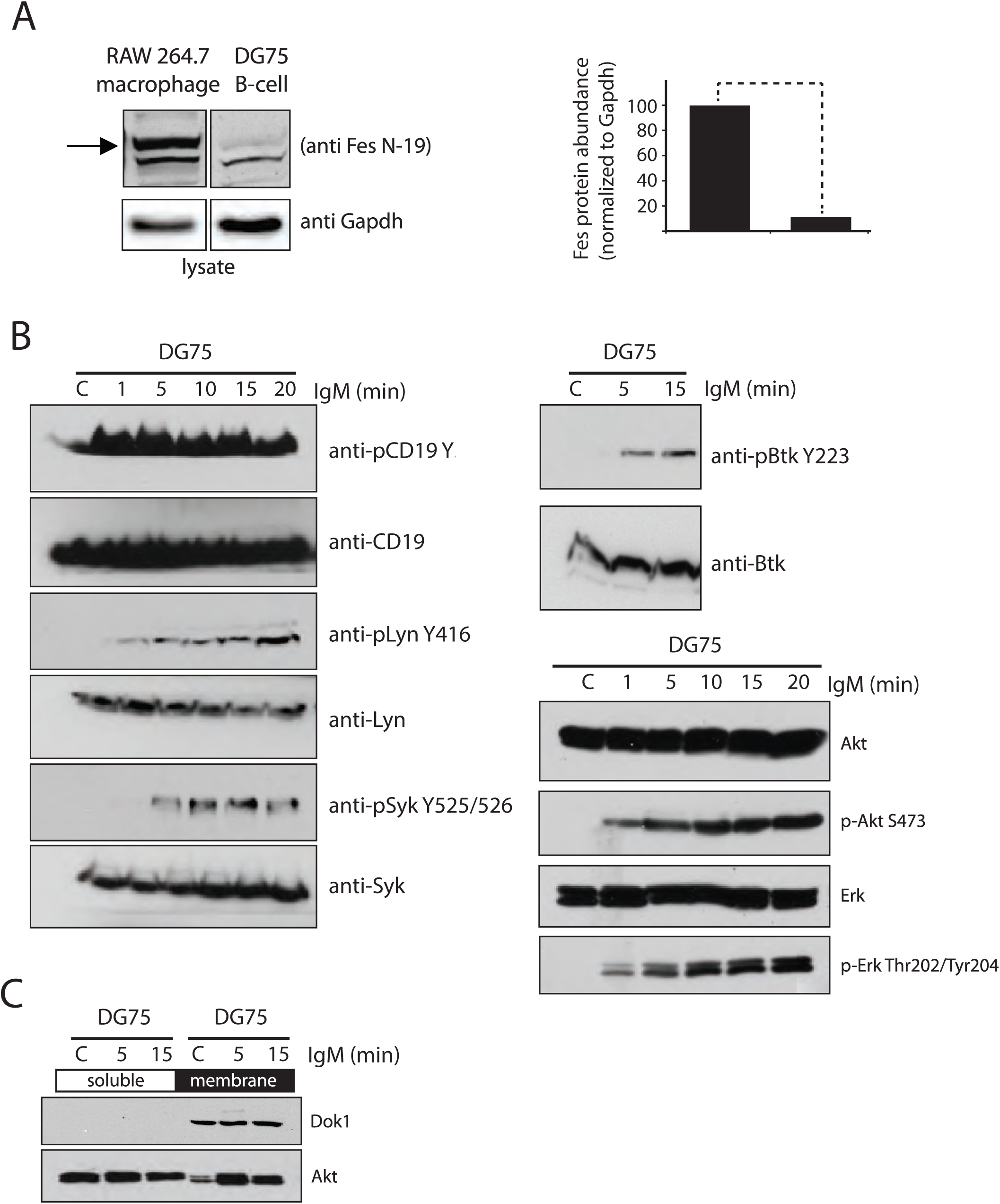
(A) Western blot analysis of endogenous Fes kinase from RAW684.3 (macrophage) and DG75 (B-lymphocyte) cell lines (Arrow indicates Fes protein band). Quantification of cellular protein levels is normalized to Gapdh and shown in the lower panel. (B) Western blot validation of activating phosphorylation sites on known BCR responders such as Syk, Btk, Lyn and CD19 showed a strong engagement of diverse signaling cascades following BCR stimulation of DG75 cells. (C) Downstream signaling pathways such as the recruitment of Akt to the membrane (Dok1 localization to the membrane is shown as a control for membrane preparations) and (D) activation of Akt and Erk pathways, as measured by phosphorylation of S473 and T202/Y204 respectively, confirmed DG75’s suitability as a model system for BCR activation and signal processing.

**Figure S2.**
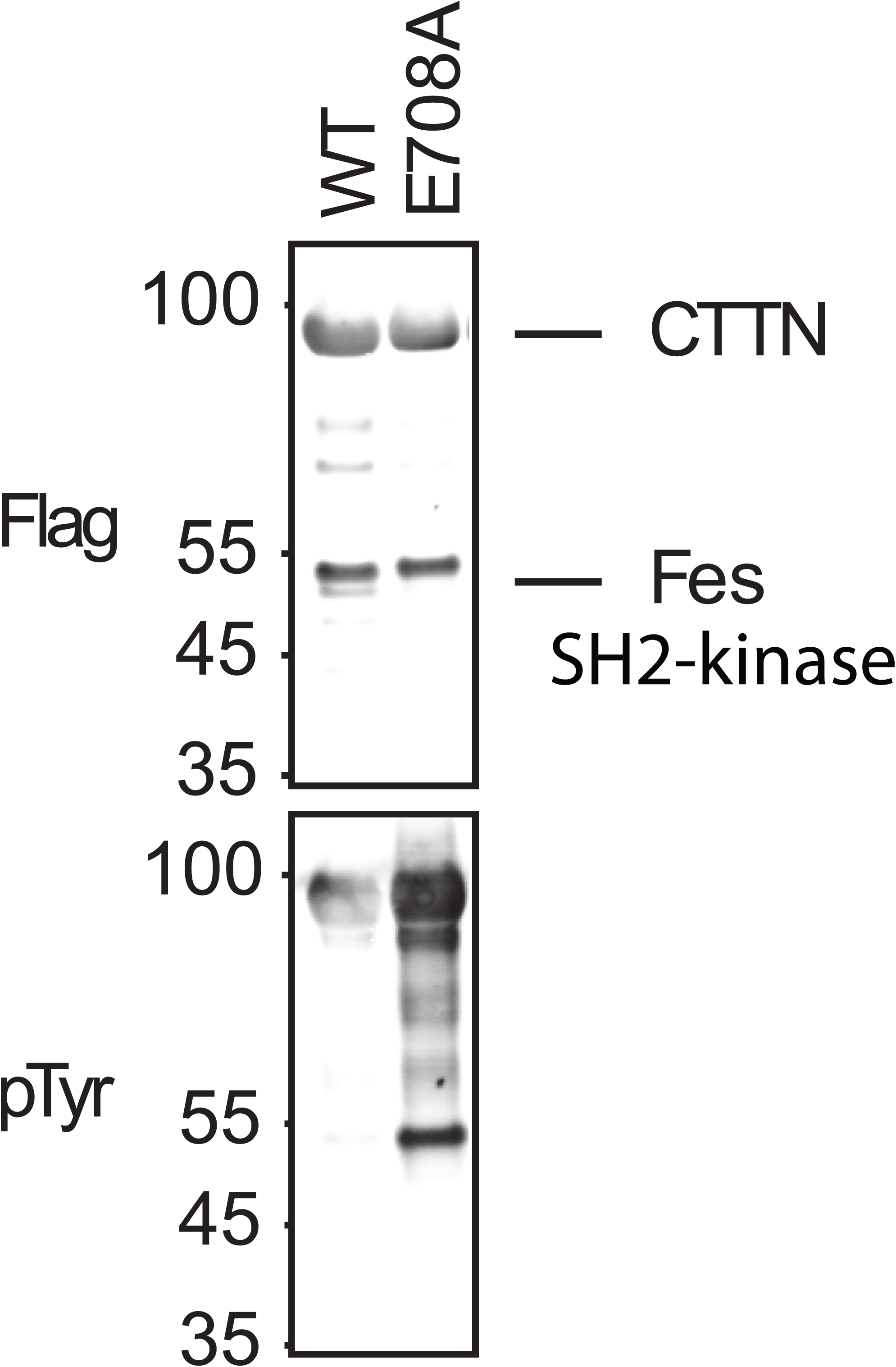
Evaluation of Cttn phosphorylation by the Fes SH2-kinase using co-expression and immunoblotting. Flag-tagged Cttn, Flag-tagged Fes SH2-kinase and Fes E708A SH2-kinase were transfected into DG75 cells and immunoprecipitated with Flag M2 antibody. Immunoblotting revealed increased autophosphorylation of Fes E708A as well as increased phosphorylation of the Cttn substrate as judged by anti-phosphotyrosine staining.

