## Supplementary figures and images for "Decoding the functional Fes kinase signaling network topology in a lymphocyte model"

### Supplemental Figure 1

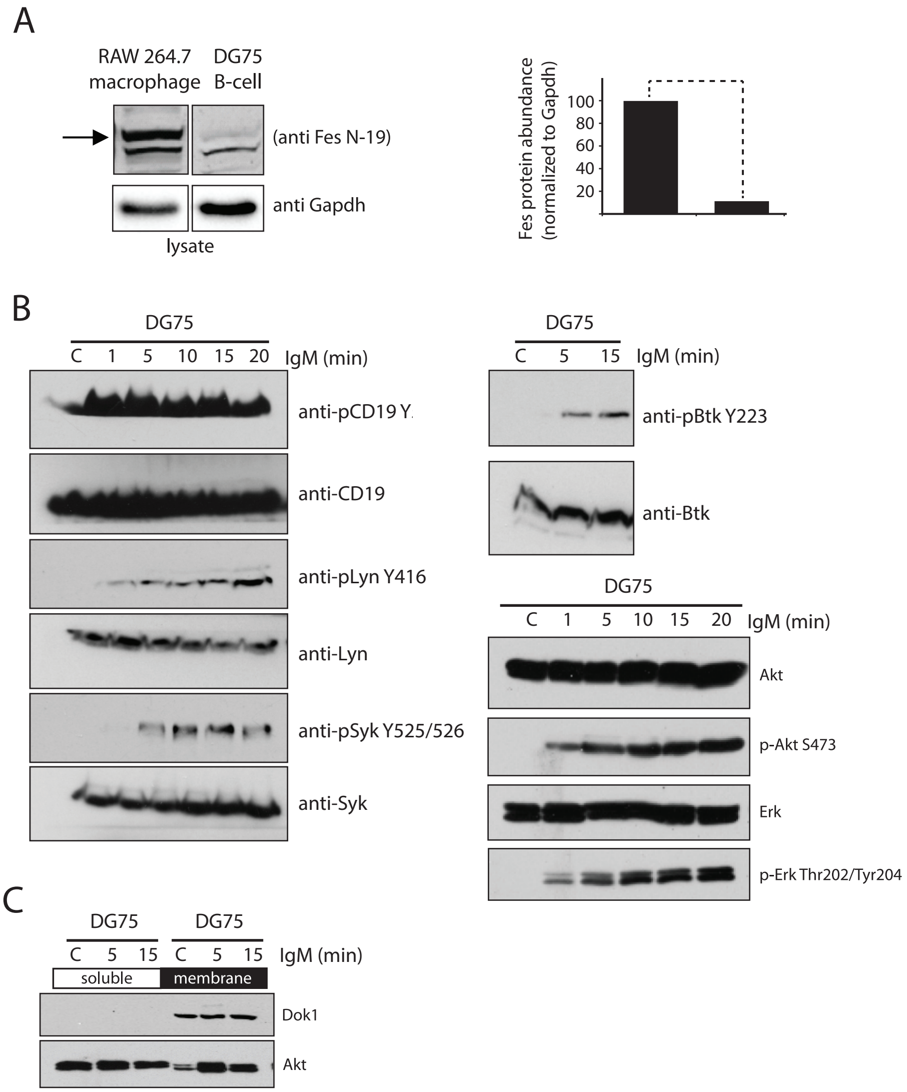

### Supplemental Figure 2

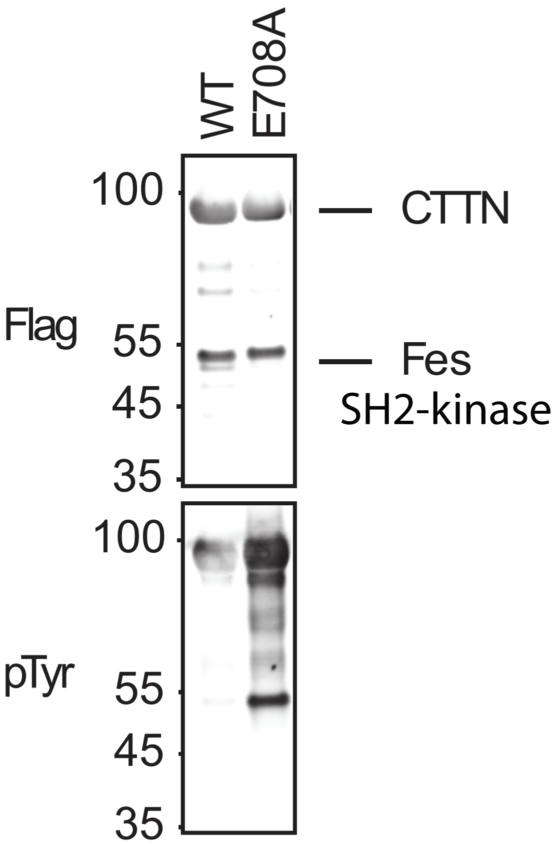
